# Human Heart Failure Alters Mitochondria and Fiber 3D Structure Triggering Metabolic Shifts

**DOI:** 10.1101/2023.11.28.569095

**Authors:** Zer Vue, Peter T. Ajayi, Kit Neikirk, Alexandria C. Murphy, Praveena Prasad, Brenita C. Jenkins, Larry Vang, Edgar Garza-Lopez, Margaret Mungai, Andrea G. Marshall, Heather K. Beasley, Mason Killion, Remi Parker, Josephs Anukodem, Kory Lavine, Olujimi Ajijola, Bret C. Mobley, Dao-Fu Dai, Vernat Exil, Annet Kirabo, Yan Ru Su, Kelsey Tomasek, Xiuqi Zhang, Celestine Wanjalla, David L. Hubert, Mark A. Phillips, Jian-qiang Shao, Melanie R. McReynolds, Brian Glancy, Antentor Hinton

**Author notes:** Corresponding Author: Antentor Hinton, Department of Molecular Physiology and Biophysics, Vanderbilt University. These authors share co-first authorship (Zer Vue, Peter T. Ajayi). These authors share senior authorship (Brian Glancy, Antentor Hinton).

## Abstract

This study, utilizing SBF-SEM, reveals structural alterations in mitochondria and myofibrils in human heart failure (HF). Mitochondria in HF show changes in structure, while myofibrils exhibit increased cross-sectional area and branching. Metabolomic and lipidomic analyses indicate concomitant dysregulation in key pathways. The findings underscore the need for personalized treatments considering individualized structural changes in HF.

## Letter Text

Mitochondria are critical in the heart, providing the energy needed for regular heart function; recent research has identified that mitochondrial dysfunction contributes to heart failure (HF), and understanding characteristics of mitochondrial dysfunction in failing hearts may assist in developing new targets for treatment^1^. In experimental models of HF, mitochondria frequently become swollen and/or fragmented, with disorganized cristae ^2^. Alteration in mitochondrial ultrastructure, may affect the efficiency of ATP production, and may account for mitochondrial dysfunction in heart failure^2^. This offers a plausible structural-dependent mechanism by which mitochondrial dysfunction in HF contributes to pathophysiology.

To explore this paradigm, we used a previously established method of serial block face-scanning electron microscopy (SBF-SEM) ^3^, to perform 3D reconstruction of mitochondria in similarly aged human samples with and without HF (Fig. A). The wide x- and y-plane dimensions of SBF-SEM make it ideal for studying mitochondrial biogenesis, networks, and alterations across regions of the heart ^3^. From the left ventricle, intermyofibrillar mitochondria, which are located between myofibrils, were manually segmented and analyzed (Fig. A). Mitochondria in HF had increased volume, surface area, and perimeter, with a tremendous inter-sample variability (Fig. B). This increased size indicates a greater capacity for ATP generation ^3^. Further consideration of how mitochondrial complexity alters shows that in HF, mitochondria take much more complex and less spherical phenotypes (Fig. C). Mitochondrial shapes including donut phenotypes and nanotunnels occur with HF, while the majority of control samples are more spherical, although each patient exhibited unique mitochondrial shapes. Thus, mitochondrial structure demonstrates tremendous variability, with certain structures that may be characteristic of HF.

From there, the myofibrillar apparatus was considered per previous techniques ^4^. The myofibrillar apparatus is the primary site of ATP utilization in the heart and plays a crucial role in subcellular remodeling including undergoing significant changes at the structural level during the development of HF. Myofibrils in HF had a greater cross-sectional area and reduced circularity per myofibril compared to control myofibrils (Fig. D). The percentage of myofibrils within the field of view of our datasets with at least one branching sarcomere, as well as the frequency of branches (Fig. D) was highest during HF as compared with control. Under control conditions, the myofibrillar apparatus was highly aligned, characterized by a prominent peak at 0° (representing perfect alignment) and another peak at 8° (Fig. D). However, in the HF condition, a greater proportion of myofibrils were observed to be oriented at angles of 0°, 7°, and 45°. Thus, cardiomyocyte sarcomere branching is a pathological feature and reflects parallel remodeling of the major ATP-producing and utilizing machinery during heart failure.

From there, we shifted our attention to how these 3D structural rearrangements concomitantly occur alongside altered metabolomics and lipidomics. As previously established ^**3**^, principal component analysis showed tremendous differences in enriched metabolites upon HF (Fig. E). Heatmaps show increased expression of numerous metabolites in HF, notably, pathway enrichment shows upregulation of signaling pathways including aminoacyl-tRNA biosynthesis and pentose phosphate pathway (PPP) (Fig. E). Notably, the PPP plays a critical role in modulating oxidative stress and glucose oxidation^**5**^, suggesting dysregulation of it may be associated with abhorrent mitochondrial function characteristic of HF. Beyond this, riboflavin, also known as vitamin B2, is a precursor for coenzymes flavin mononucleotide and flavin adenine dinucleotide, which is important in energy production ^**6**^. Through co-activation of acyl-CoA, riboflavin may act in a compensatory mechanism, reducing oxidative stress and alliterating impaired energy production ^**6**^. Lipidomic analysis further shows that although lipid length did not change, there are differences among individuals with heart failure with upregulation of classes including acylcarnitines (Fig. F). These have previously arisen as key biomarkers in HF ^**7**^. Together, this illustrates that mitochondrial and fiber structural alterations concurrently occur alongside altered enrichment pathways, indicating potential mechanisms to restore pathological structure.

Given that mitochondrial bioenergetics are necessitated in both systolic and diastolic heart function ^1,2^, the mechanisms that govern and alter mitochondria in HF cases remain an intriguing future avenue. Past research has found that in HF, mitochondria exhibit key signs of dysfunction including decreased ETC activity, changes in ion activity, and altered dynamics ^1,2^. Here, we have also established that varying subpopulations of mitochondria undergo changes in nanotunnels and mitochondrial structural arrangement, while the wider spatial orientation of cardiac myofibrils further changes. Our results highlight the importance of consideration of mitochondria structure in the treatment of HF, which may vary tremendously between individuals, both dependent and independent of HF status. Broadening physiological implications of these unique shapes, and how their relative abundance may be affected by protein quantity and spatial arrangement may offer insight for personalized medicine. While past studies have looked at general age-dependent mitochondrial structural changes ^**3**^, equally important is the consideration of mitochondrial remodeling that occurs in an age-dependent manner after challenges including HF. Other types of HF may further display different mitochondrial organization, metabolism, and lipid distribution which must be further explicated. Consideration of how mitochondria and myofibril structure change in dependence on pathophysiology, may offer an avenue for individualized medicine that targets HF through modulation of mitochondrial structure.

## Figure and Legend

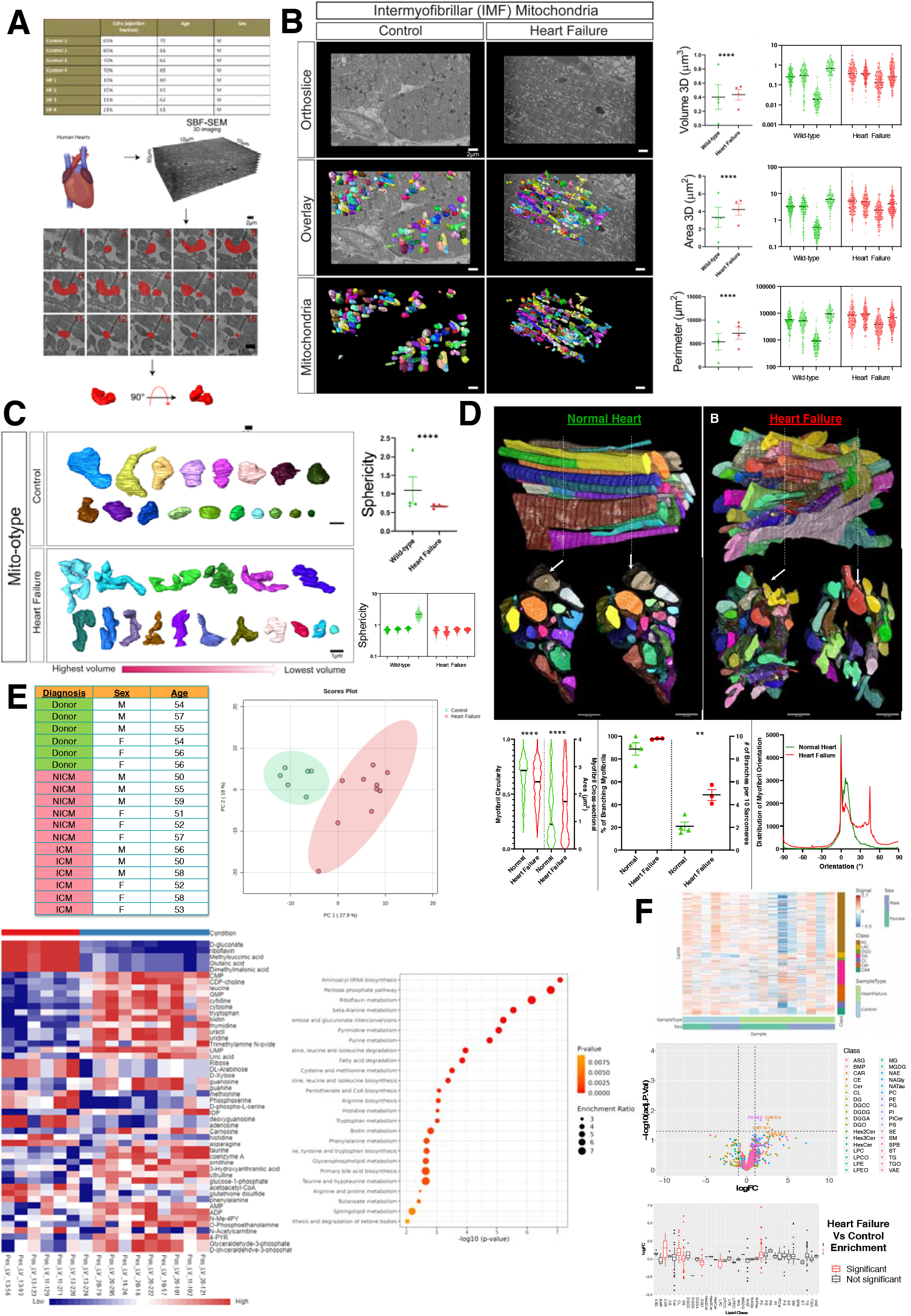
(**A**) Through existing institutional review board approval (#41148), male human heart failure (HF) dilated cardiomyopathy samples (ejection fraction between 10-21% and ages between 60-63) and controls (ejection fraction between 69-80% and ages between 62-81) were collected (n=4). Serial block face scanning electron microscopy (SBF-SEM) was utilized for manual contour segmentation of mitochondria from orthoslices. (**B**) Representative SBF-SEM orthoslice, 3D reconstructions of mitochondria, and isolated mitochondria in control and HF. Mitochondrial volume, surface area, and perimeter are all displayed, with the averages of each sample shown to the left, while the right shows dots representing values of each mitochondrion in samples. (**C**) Mito-otyping, a method of organizing mitochondria on the basis of their volume, shows the relative changes in mitochondria structure in HF samples, which are quantified based on sphericity. (**D**) Raw SBF-SEM image volumes were rotated in 3D to visualize the muscle cell’s cross-section. Myofibrillar cross-sectional area (CSA) and circularity were measured by converting the traced structures to binary images and using the Analyze Particles plugin in ImageJ for each slice throughout the volume (control n□=□4 humans, 6 cells, 133 myofibrils, 1118 sarcomeres; HF n□=□3 humans, 12 cells, 219 myofibrils, 2428 sarcomeres). The distribution data reflects how much of the myofibrillar volume is perfectly aligned (0°) versus misaligned (away from 0°). (**E**) Metabolomic analysis comparing Ischemic Cardiomyopathy (ICM), Non-Ischemic Cardiomyopathy (NICM), and donor samples (n=6) from a mixture of age-matched males and females. Principal component analysis and metabolic heatmap showing the relative abundance of metabolites in control and HF. Enrichment analysis and pathway impact for metabolites enriched in HF. (**F**) Using the same samples, lipidomic analysis compared HF and control, showing differences in lipid classes by sex in heatmaps, lipid class (as shown in volcano plots and box plot), and differences in lipid chain length. For all panels, dot-plots show mean±SEM, and the numbers of independent samples are indicated, ^****^, *p* < 0.0001; ^***^, *p* < 0.001; ^**^, *p* < 0.01; ^*^, *p* < 0.05, calculated with Student’s *t-*test.

## FUNDING

This project was funded by the National Institute of Health (NIH) NIDDK T-32, number DK007563 entitled Multidisciplinary Training in Molecular Endocrinology to Z.V.; National Institute of Health (NIH) NIDDK T-32, number DK007563 entitled Multidisciplinary Training in Molecular Endocrinology to A.C.; Integrated Training in Engineering and Diabetes”, Grant Number T32 DK101003; Burroughs Wellcome Fund Postdoctoral Enrichment Program #1022355 to D.S.; The UNCF/ Bristol-Myers Squibb (UNCF/BMS)-E.E. Just Postgraduate Fellowship in Life sciences Fellowship and Burroughs Wellcome Fund/ PDEP #1022376 to H.K.B.; NSF MCB #2011577I to S.A.M.; NSF EES2112556, NSF EES1817282, NSF MCB1955975, and CZI Science Diversity Leadership grant number 2022-253614 from the Chan Zuckerberg Initiative DAF, an advised fund of Silicon Valley Community Foundation to S.D.; The UNCF/Bristol-Myers Squibb E.E. Just Faculty Fund, Career Award at the Scientific Interface (CASI Award) from Burroughs Welcome Fund (BWF) ID # 1021868.01, BWF Ad-hoc Award, NIH Small Research Pilot Subaward to 5R25HL106365-12 from the National Institutes of Health PRIDE Program, DK020593, Vanderbilt Diabetes and Research Training Center for DRTC Alzheimer’s Disease Pilot & Feasibility Program. CZI Science Diversity Leadership grant number 2022-253529 from the Chan Zuckerberg Initiative DAF, an advised fund of Silicon Valley Community Foundation to A.H.J.; and National Institutes of Health grant HD090061 and the Department of Veterans Affairs Office of Research award I01 BX005352 (to J.G.). Howard Hughes Medical Institute Hanna H. Gray Fellows Program Faculty Phase (Grant# GT15655 awarded to M.R.M); and Burroughs Wellcome Fund PDEP Transition to Faculty (Grant# 1022604 awarded to M.R.M). Additional support was provided by the Vanderbilt Institute for Clinical and Translational Research program supported by the National Center for Research Resources, Grant UL1 RR024975–01, and the National Center for Advancing Translational Sciences, Grant 2 UL1 TR000445–06 and the Cell Imaging Shared Resource. Its contents are solely the responsibility of the authors and do not necessarily represent the official view of the NIH. The funders had no role in study design, data collection and analysis, decision to publish, or preparation of the manuscript.

## CONFLICT OF INTEREST

The authors declare that they have no conflict of interest.

## Data Availability

The methods, data, and materials are available upon request.

## Notes

### Competing Interest Statement

The authors have declared no competing interest.

### Summary of Updates

Author list update

